# Deficiency of m^6^A RNA methylation promotes ZBP1-mediated cell death

**DOI:** 10.1101/2024.06.29.601251

**Authors:** Shuang Li, Xiangyu Deng, Deepak Pathak, Rashmi Basavaraj, Lina Sun, Yating Cheng, Jian-Rong Li, Marissa Burke, Gavin W. Britz, Chao Cheng, Yang Gao, Yi-Lan Weng

## Abstract

m^6^A RNA methylation suppresses the immunostimulatory potential of endogenous RNA. Deficiency of m^6^A provokes inflammatory responses and cell death, but the underlying mechanisms remain elusive. Here we showed that the noncoding RNA 7SK gains immunostimulatory potential upon m^6^A depletion and subsequently activates the RIG-I/MAVS axis to spark interferon (IFN) signaling cascades. Concomitant excess of IFN and m^6^A deficiency synergistically facilitate the formation of RNA G-quadruplexes (rG4) to promote ZBP1-mediated necroptotic cell death. Collectively, our findings delineate a hitherto uncharacterized mechanism that links m^6^A dysregulation with ZBP1 activity in triggering inflammatory cell death.

## Introduction

The innate immune system uses sophisticated surveillance mechanisms that adeptly distinguish foreign DNA/RNA among host nucleic acids(1,2). Such self/non-self discrimination establishes appropriate immune defenses against fungal, bacterial, and viral pathogens, but prevents spontaneous reactions to endogenous nucleic acids. However, under certain physiological and pathological conditions, endogenous RNA may be misprocessed to acquire immunostimulatory potential(3). These host-derived immunostimulatory RNA manifest structural characteristics analogous to viral double-stranded RNA (dsRNA), potentially confounding the host’s pattern recognition receptors (PRRs) and erroneously activating antiviral pathways even in the absence of an actual viral infection. This phenomenon, termed the “viral mimicry response”(4), prompts the activation of interferon (IFN) signaling, leading to unwarranted immune responses and inflammatory cell death.

RNA modifications represent one of such self/non-self discrimination mechanisms that chemically annotate the host RNAs, thereby attenuating their innate immunogenicity(5). Among 150 distinct chemical RNA modifications, several studies have elucidated that adenosine-to-inosine (A-to-I) editing within inverted-repeat Alu elements (IR-Alus) can perturb RNA duplex conformation, consequently attenuating the immunogenic potential of these specific RNA species(6,7). Despite recent advancements in transcriptome-wide elucidation of RNA modifications, whether and how these modifications, aside from A-to-I editing, regulate the emergence of endogenous immunostimulatory RNA remains elusive.

m^6^A is a prevalent modification commonly found in messenger RNAs (mRNAs) and long non-coding RNAs. This modification, established by the METTL3/METTL14 methyltransferase complex, is well-recognized to influence diverse aspects of RNA metabolism, including RNA stability and translation. Intriguingly, recent findings indicate that perturbations of m^6^A can induce a viral mimicry response(8–10), underscoring the potential role of m^6^A in suppressing host RNA immunogenicity. However, the specific host RNA entities that acquire immunostimulatory characteristics upon m^6^A perturbation remain inadequately understood.

IFN signaling activates the expression of myriad interferon-stimulated genes (ISGs) with a wide range of immunomodulatory functions. Overamplification of IFN signaling can lead to cell death, which can result in potential tissue degeneration and increased mortality. Emerging evidence suggests that Z-DNA-binding protein 1 (ZBP1), an ISG, may interact with enigmatic Z-RNA/DNA, subsequently activating cell death cascades(11–13). Given recent findings indicating m^6^A deficiency adversely affects cellular survival, we postulate that m^6^A deficiency may potentiate the interactions between ZBP1 and its cognate RNA ligands, thereby leading to cellular death and tissue degeneration.

In this study, we report that m^6^A deficiency activates the RIG-I/MDA5 signaling pathway, prompting a viral mimicry response. We further unveil that 7SK snRNA (RN7SK), a noncoding RNA with a 5’-ppp moiety, acquires immunogenic properties in the absence of m6A RNA methylation. Using bioinformatics and cellular analyses, we demonstrate that m^6^A deficiency or IFN induction promotes RNA G-quadruplexes (rG4s) formation, thereby facilitating ZBP1 interaction. These findings reveal that m^6^A reduces the immune reactivity of certain host RNA subsets and offer new insights into how m^6^A dysregulation links to ZBP1 mediated cell death and outlines a potential pathological mechanism in related disorders.

## Materials and Methods

### Generation of METTL3 and YTHDC1 Knockout (KO) cell lines

Single guide RNA (sgRNA) sequences targeting METTL3 or YTHDC1 were designed using Benchling (https://www.benchling.com/). Detailed sgRNA sequences are provided in the Supplementary Table 5. The chosen sgRNA for METTL3 or YTHDC1 was cloned into the pCAG-eCas9-GFP-U6-gRNA plasmid (Addgene #79145), which harbored the green fluorescent protein (GFP) marker for visual identification of transfected cells. HMC3 cells were transfected with this plasmid using Turbofect transfection reagent. After 48 hours of post-transfection, cells with GFP signal were isolated using fluorescence activated cell sorting (FACS) and individually sorted into separate wells of a 96-well plate. The resulting single-cell colonies underwent screening using Sanger sequencing and Western blot analysis to confirm the successful knockout of METTL3.

For establishing METTL3 KO cell lines with an additional gene knockout, previously established stable METTL3 KO cells underwent transfection with the pCAG-eCas9-GFP-U6-gRNA vector, this time carrying a different sgRNA. Cells exhibiting double knockouts were identified and selected based on their GFP expression, after which they were further cultured for use in subsequent experimental assays.

### Generation of ZBP1 overexpressing stable cell lines

To generate cell lines stably overexpressing ZBP1, the full-length ZBP1 gene was cloned into the PGAMA lentiviral expression vector (Addgene #74755). This vector co-expresses mCherry, allowing easy identification of successfully transduced cells. Lentiviral particles were produced by co-transfecting the PGAMA-ZBP1 construct, along with packaging plasmids psPAX2 and pMD2G, into HEK 293T cells using a transfection ratio of 4:3:1. Approximately 48 hours post-transfection, the viral supernatant was harvested and subjected to centrifugation to remove any cellular debris and intact cells.

This cleared viral supernatant was subsequently used to transduce both wild-type (WT) and METTL3 KO cells that had been previously plated in 6-well culture plates. After 7 days of incubation to allow for viral integration, cells displaying mCherry fluorescence were isolated using FACS. These mCherry-positive cells were then pooled to form stable cell lines with consistent overexpression of ZBP1.

### RNA extraction, RT-qPCR analysis, and library construction

Total RNA was extracted using TRIzol reagent (Invitrogen) according to the manufacturer’s instructions. The RNA was then purified and concentrated using RNA Clean and Concentrator-5 spin columns (Zymo Research; Catalog #R1015).

For the generation of mRNA sequencing libraries, we employed the SMART-seq protocol as previously described(14). Three biological replicates were prepared for each experimental condition and uniquely barcoded. These barcoded samples were pooled in equimolar concentrations and sequenced on an Illumina NextSeq 500.

For constructing ribosomal RNA (rRNA)-depleted RNA-seq libraries, 100 ng of total RNA was first subjected to rRNA removal using the NEBNext rRNA Depletion Kit (Human/Mouse/Rat). The rRNA-depleted RNA was then processed using the NEBNext Ultra II Directional RNA Library Prep Kit for Illumina, following the manufacturer’s protocol.

For RT-qPCR, cDNA was synthesized from 1 μg of extracted RNA per sample using the SMARTScribe reverse transcriptase (Clontech) and dT primer (GTCTCGTGGGCTCGGAGATGTGTATAAGAGACAG T30VN). The RT-qPCR reactions were conducted using SYBR Green Master Mix (Roche) on a C1000 Touch Thermocycler equipped with a CFX96 Real-Time System (Bio-Rad). Gene expression levels were normalized to housekeeping genes, GAPDH or Tubulin, as internal controls. All primers utilized in the RT-qPCR are provided in the Supplementary Table 5.

### Immunostimulatory RNA assay

Total RNA was extracted from both WT and METTL3 KO cells using Trizol reagent, followed by purification with RNA Clean and Concentrator-5 spin columns. To isolate specific RNA fractions, rRNA was removed from the total RNA using the NEBNext rRNA Depletion Kit (Human/Mouse/Rat). For targeted isolation of RNAs of interest, such as 7SK or 7SL, biotinylated antisense oligonucleotides (ASOs) were employed. A mixture containing 20 µg of total RNA and 2 µg of ASO in 200 µl of RNA binding buffer (20 mM Tris-HCl, pH 7.5, 1.0 M LiCl, and 2 mM EDTA) was incubated at 70°C for 5 minutes, followed by a 10-minute incubation at room temperature. The RNA-ASO mixture was then combined with streptavidin magnetic beads (Thermo Fisher) and agitated continuously for 30 minutes at 25°C. Subsequent washes (three times) with RNA wash buffer (10 mM Tris-HCl, pH 7.5, 0.15 M LiCl, and 1 mM EDTA) were performed to remove non-specifically bound RNA. The specifically bound target RNA was then eluted using Trizol and further purified using RNA Clean and Concentrator-5 spin columns.

For the functional assays, equal quantities of RNA from WT and METTL3 KO cells were transfected into WT cells using Lipofectamine RNAiMAX. At 24 hours post-transfection, cells were lysed with 300 µL of TRIzol for subsequent RNA extraction. RT-qPCR analysis was used to measure the induction of ISGs, assessing the immunostimulatory properties of the RNA samples.

### Immunofluorescence staining

Cells seeded on coverslips were first fixed with 4% paraformaldehyde (PFA) in PBS at room temperature for 15 minutes. They were then permeabilized using 0.1% Triton X-100 in PBS (PBST) for another 15 minutes and blocked with 3% normal donkey serum for an hour. The coverslips were then incubated with primary antibodies overnight at 4°C. After washing four times with PBST, the samples were incubated with secondary antibodies (1:300) and DAPI (1:5000) for 2 hours at room temperature. After washing four times with PBST, the coverslips were mounted, and images were acquired using a confocal microscope.

To detect and quantify apoptotic or dying cells, we utilized YOYO-1 staining as previously described(12), staining nucleic acids in cells with compromised membrane integrity. In brief, cells were treated with 10 nM YOYO-1 iodide dye and incubated for 30 minutes, allowing the dye to bind to exposed nucleic acids in cells with damaged membranes. Following this, cells were washed to remove unbound dye, then fixed with 4% PFA and counterstained with DAPI for imaging.

### Western blot analysis

Proteins extracted from WT, METTL3 KO, and YTHDC1 KO cells were resolved on 10% Mini-PROTEAN TGX Precast Protein Gels (Bio-rad) and subsequently transferred to a PVDF membrane. This membrane was blocked using 5% dry milk for an hour at room temperature, followed the incubation of the anti-METTL3 antibody (Abcam; 1:1000) or anti-YTHDC1 antibody (Proteintech, 1:1000) overnight at 4°C. After thorough washing, the membrane was probed with the secondary HRP-conjugated anti-rabbit IgG antibody (Santa Cruz; 1:10000). To ensure equal protein loading, the membrane was also probed with mouse anti-GAPDH (EMD Millipore; AB2302).

### Cell viability assays

For crystal violet staining, HMC3 cells that underwent the designated genetic manipulation were seeded in six-well plates at a density of 0.8 x 10^5^ cells per well. After 72-hour growth, the cells were fixed with a 4% paraformaldehyde and subsequently stained with a 1% crystal violet solution for 10 minutes. For assessing cell viability, images were captured from five distinct fields within each well. Cell growth was then evaluated by measuring the area of the stained cells.

For interferon treatment assays, HMC3 cells were initially seeded at a density of 2000 cells per well in 96-well plates, each containing 100 µl of culture medium. The following day, cells were treated with IFNb (200ng/mL) or PBS (vehicle). The cells were cultured for a duration of 0 to 5 days. Subsequently, 20 µl of the CellTiter-Blue reagent was directly added to each well, and the plates were incubated for 2 hours at 37°C. Fluorescence measurements were recorded with excitation at 560 nm and emission at 590 nm. This assay was performed at 24-hour intervals to continuously monitor cell viability for 5 days.

### Bimolecular fluorescence complementation (BiFC) assay

The gateway cloning technique was employed to incorporate the ZBP1 gene into GFP10C and GFP11C vectors, resulting in the formation of two ZBP1 constructs, each possessing complementary fragments of GFP. These constructs were used to transfect cells using the Turbofect transfection reagent, enabling the evaluation of ZBP1 oligomerization based on the reconstitution of the split GFP fragments. Following transfection, the cells were treated with IFNb (100ng/mL) or with STM2457, a METTL3 inhibitor (10 µM), for a designed duration. After the treatment, the cells were fixed with 4% PFA and counterstained with DAPI in preparation for imaging.

### RNA immunoprecipitation (RIP)-qPCR

For RIG-I RIP-qPCR, cells were lysed on ice for 10 minutes using 500 µl of lysis buffer (20 mM Tris-Cl, pH 7.4, 150 mM NaCl, 5 mM MgCl2, 1 mM DTT, 0.5% NP-40) supplemented with 5 µl of SUPERase•In RNase Inhibitor (Invitrogen). This lysate was centrifuged at maximum speed for 15 minutes at 4°C, and the clear soluble extract was collected. The extract was then divided into two equal parts. One portion was incubated with 2 µg of anti-RIG-I antibody and the other with 2 µg of Rabbit IgG, both at 4°C for 2 hours. Pre-washed Dynal protein A/G magnetic beads (Invitrogen) in lysis buffer were then added to the soluble extracts and incubated for an additional hour at 4°C. The beads were washed four times using the lysis buffer. RNA from the beads was extracted using TRIzol and then purified using RNA Clean and Concentrator-5 spin columns.

The isolated RNAs were subsequently reverse transcribed using random hexamer primers and SMARTScribe Reverse Transcriptase (Clontech; 639537). Indicated genes were analyzed by RT-qPCR using Fast SYBR Green Master Mix (ThermoFisher; 4385612). The sequences of the primers utilized for each gene are listed in the Supplementary Table 5.

### RNA sequencing data analysis

After removing low-quality reads, the remaining high-quality sequences were aligned to the human genome reference (GRCh38) using the STAR aligner. Differential gene expression analysis was then conducted using the DESeq2 package. For this study, genes exhibiting absolute fold change of ≥1.5 and adjusted p-value (p-adj) <0.05 were categorized as significantly differentially expressed genes (DEGs).

### ADAR1 RIP-seq analysis

To discern RNA transcripts enriched in the ADAR1 P110 and ADAR1 P150 RIP-seq assays, comparative analyses of RNA expressions from each antibody immunoprecipitation against their corresponding input RNA levels were conducted using the DESeq2 package. Transcripts exhibiting fold change of ≥1.5 and p-adj of <0.05 were identified as significantly DEGs. The genomic distribution of the reads obtained from the ADAR1 P110 and ADAR1 P150 RIP-seq was further analyzed using the RSeQC package. For comparative purposes, rG4 transcripts identified in prior studies were employed in downstream analyses.

### A-to-I RNA editing analysis

The identification of high-confidence A-to-I RNA editing sites was performed as previously described(15). Briefly, ADAR1 110 and 150 RIP-seq datasets were aligned to the human genome reference (GRCh38) using the STAR aligner. A-to-G mismatches, indicative of A-to-I editing events, were identified through pileup2var. To qualify as high-confidence editing sites, several stringent criteria were imposed: (1) minimum of 10 mapped reads at the site, (2) at least 2 A-to-G variant occurrences, (3) an editing ratio threshold set at a minimum of 5%, and (4) the effective signal of A-to-G conversion must exceed 95%. Upon identifying these sites, bedtools was employed to aggregate editing sites into clusters. Each site within a cluster was defined as being within 100 base pairs of another editing site. ANNOVAR software was subsequently utilized for comprehensive annotation, detailing the specific genomic locations and the associated genes.

### In vitro A-to-I editing assay

Human ADAR1 P150 (amino acids 127-1226, encompassing the entire Zα domain) was cloned into the pLEX vector with a cleavable N-terminal MBP tag. ADAR1 P150 was overexpressed through transient transfection into suspension HEK293T cells and purified using Amylose resin followed by Superose S6 gel filtration.

The in vitro editing assay was conducted by incubating 700 nM ADAR1 P150 with 2 μM EIF4EBP2 RNA in a buffer containing 10 mM HEPES, 70 mM KCl, 5% Glycerol, and 1 mM DTT at 37°C for 1 hour, and then was halted by heating at 85°C for 5 minutes. For gel assays assessing EIF4EBP2 editing, the reaction was further incubated with 100 nM EcEndoV (homemade) at 37°C for 45 minutes, then terminated by adding 2x loading buffer (93.5% formamide, 0.025% xylene cyanol FF, and 50 mM EDTA, pH 8.0) and heated at 85°C for 5 minutes. EcEndoV preferentially recognizes and cleaves modified inosines within RNA, generating shorter RNA fragments for gel electrophoresis detection. The quenched reactions were resolved on 12% denaturing polyacrylamide gels (Urea-PAGE) and stained with SYBR Gold. To determine the editing percentage of EIF4EBP2, RNA was purified post-ADAR1 P150 editing using an RNA cleanup kit, reverse transcribed into cDNA, and subsequently subjected to Sanger sequencing. Editing events (A-to-G mutations) were estimated by comparing the relative peak areas of guanine versus adenine in the ABI sequencing trace.

### ZBP1 STAMP analysis

The Zαβ domain of human ZBP1 was cloned into the pLIX403_Capture1_APOBEC vector (Addgene #183951) utilizing the gateway cloning method. The resulting construct, pLIX403_ZBP1zαβ_APOBEC (ZBP1zαβ_APOBEC), was transfected into HEK293 cells.

Doxycycline (dox) was administered 24 hours post-transfection to induce protein expression. RNA samples were collected 24 hours after induction and subsequently analyzed via RNA-seq. The plasmid, pLIX403_APOBEC (Addgene #183901), was utilized as control experiment (APOBEC1).

High-confidence C-to-U RNA editing sites were determined as previously described(16). Briefly, sequencing reads from ZBP1zαβ-APOBEC1 and the control APOBEC1 RNA-seq data were aligned to the human genome reference (GRCh38) using the STAR aligner. C-to-U mismatches, indicative of APOBEC-mediated base conversion, were identified through pileup2var. High-confidence editing sites were then determined based on several stringent criteria: (1) a minimum of 10 mapped reads at the site, (2) at least 2 occurrences of C-to-U variants, (3) an editing ratio threshold set at a minimum of 5%, and (4) the effective signal of C-to-U conversion exceeding 95%. We then compared these sites from ZBP1zαβ-APOBEC1 with those from APOBEC1 to have ZBP1zαβ-APOBEC1 specific editing sites. High-confidence ZBP1 binding loci were determined by using bedtools to aggregate the ZBP1zαβ-APOBEC1 editing sites into clusters, with each site located within 100 bases of another.

For the enrichment analysis, we used Fisher’s exact test to compare high-confidence ZBP1 binding loci with RNA-binding protein (RBP) binding sites from eCLIP data of the ENCODE project and published YTH protein PAR-CLIP datasets(17,18). The odds ratio of Fisher’s exact test was calculated to determine the extent of enrichment.

### Detection and quantification of rG4 formation

We employed DsiRNAs (IDT) to silence the expression of YTHDF1 and YTHDF2 for 48 hours (Supplementary Table 5). Subsequently, we assessed the formation of RNA G-quadruplexes (rG4) using a hemin-based biotin assay, as previously described(19). Briefly, cells were treated with 100 µM hemin for 30 minutes, then incubated with 100 µM biotin phenol for 5 minutes.

Afterwards, cells were exposed to 100 µM H2O2 for one minute to quench the reaction. Post fixation with PFA, cells were treated with Cy3-conjugated streptavidin antibody and imaged to gauge the level of biotinylation.

### Quantification and statistical analysis

Data visualization and statistical analyses were conducted using R and GraphPad Prism. For comparing two groups, the unpaired 2-tailed t-test was employed for samples with a normal distribution, and the Mann-Whitney test was used for those with a non-normal distribution. For evaluations involving more than two groups, a one-way ANOVA followed by Tukey’s post hoc test was utilized. Significance levels are denoted as: *P<0.05 and **P<0.01.

## Results

### m^6^A deficiency induces viral mimicry responses

In this study, we employed CRISPR/Cas9 technology to generate METTL3 knockout (KO) in primary derived microglia (HMC3) to investigate the role of m^6^A RNA methylation in regulating microglial activation and cellular responses. Western blot analysis confirmed the absence of METTL3 expression in the KO cell lines (Figure 1A). Notably, METTL3 KO cells displayed significant growth defects (Figure 1B). To elucidate the molecular mechanisms through which m^6^A deficiency impacts cell vitality, we performed RNA-seq analysis on METTL3 KO and wild-type (WT) cells. Our analysis revealed 736 upregulated and 698 downregulated genes in the METTL3 KO microglia (Figure 1C and Supplementary Table 1). Gene Ontology (GO) analysis revealed a significant enrichment in pathways related to innate immune and antiviral responses among the upregulated genes (Figure 1D). This finding supports previous studies showing that m^6^A deficiency can trigger inflammatory responses(9,10). Further transcription factor (TF) enrichment analysis highlighted an increased presence of Interferon Response Factors (IRFs) and Interferon-Sensitive Response Elements (ISREs) in upregulated genes (Figure 1E).

**Figure 1.**
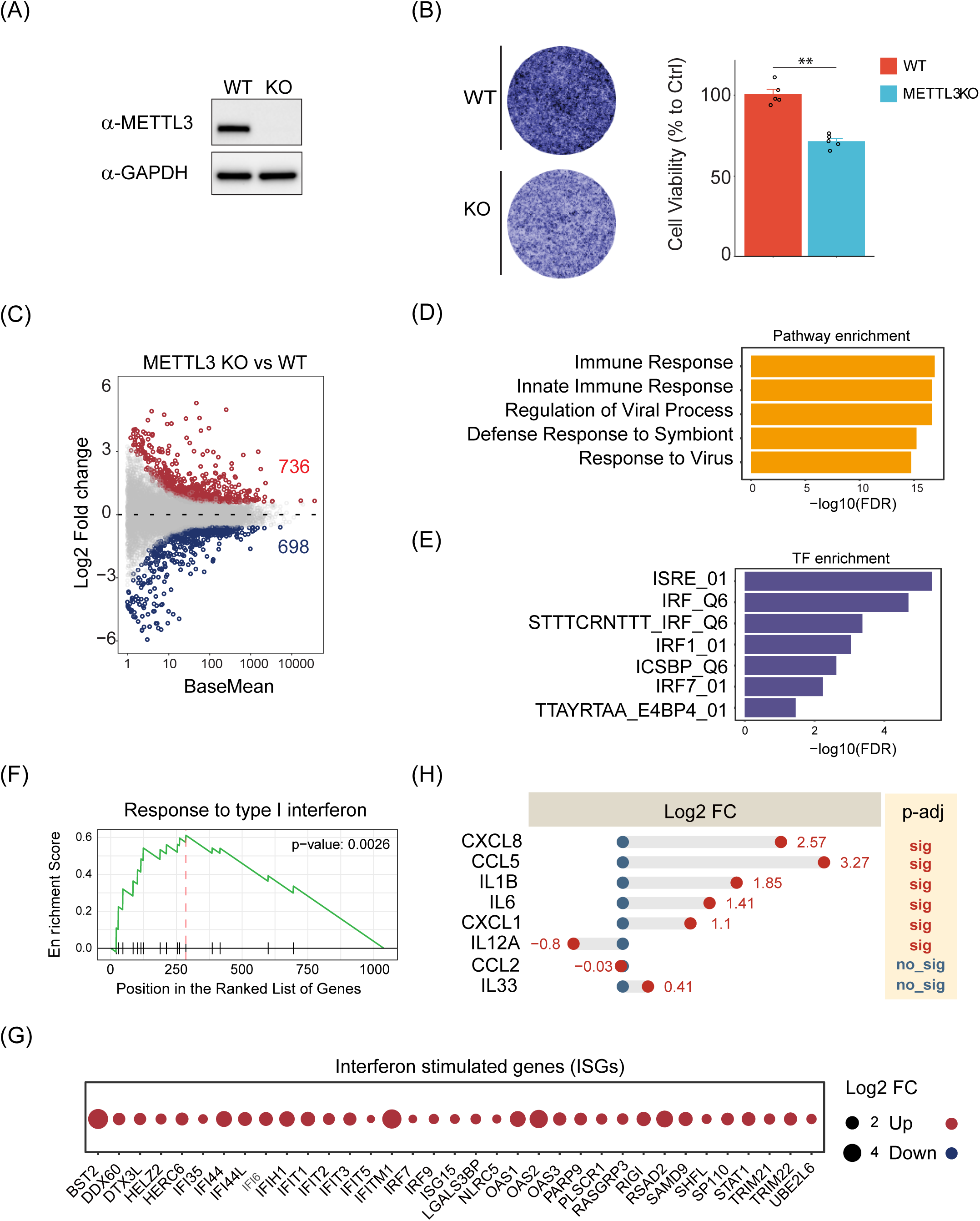
m^6^A deficiency induces viral mimicry responses. (A) Confirmation of METTL3 KO via western blot analysis. (B) Comparison of cell growth rates between WT and METTL3 KO cells, determined using crystal violet assays. Data are normalized to WT and presented as mean ± SEM (n = 5 experimental replicates; **P < 0.01, t-test). (C) RNA-seq analysis highlighting differentially expressed genes between METTL3 KO and WT cells. (D) GO analysis illustrating the overrepresentation of pathways related to innate immune responses among upregulated genes in METTL3 KO cells. (E) Enrichment analysis illustrating the top transcription factors (TFs) associated with genes that are upregulated in METTL3 KO cells. (F) Gene Set Enrichment Analysis (GSEA) showing significant enrichment in the interferon response pathway. (G) Expression levels of ISGs in METTL3 KO cells, as identified by RNA-seq. Expression values are normalized to WT and presented as log2 fold change. (H) Induction of cytokines and chemokines in METTL3 KO cells, as determined by RNA-seq. Expression levels are normalized to WT and presented as log2 fold change.

Additionally, Gene Set Enrichment Analysis (GSEA) revealed that m^6^A deficiency activates Type I interferon signaling (Figure 1F). In line with these findings, METTL3 KO cells demonstrated enhanced expression of numerous Interferon-Stimulated Genes (ISGs) (Figure 1G), along with a broad array of cytokine and chemokine activation (Figure 1H). Together, these results suggest that m^6^A deficiency contributes to a state of heightened inflammatory response activation.

m^6^A is recognized for its role in regulating RNA stability. One hypothesis for the observed upregulation of ISGs in METTL3 KO cells could be the increased RNA stability resulting from m^6^A depletion. However, our analysis of METTL3 KO and m^6^A MeRIP-seq datasets(20) does not support this hypothesis, as many m^6^A-methylated transcripts did not exhibit increased expression levels in METTL3 KO cells (Supplementary Figure S1A). Additionally, we observed no substantial induction of interferon receptors that would otherwise enhance the activation of interferon signaling pathways (Supplementary Figure S1B). Taken together, these findings suggest that the absence of m^6^A may activate inflammatory responses through alternative regulatory mechanisms rather than through changes in RNA stability of inflammatory response regulators and genes.

### m^6^A deficiency activates the RIG-I/MAVS pathway to elicit inflammatory responses

Previous research has shown that m^6^A depletion can result in an accumulation of endogenous double-stranded RNA (dsRNA) (10). It was hypothesized that innate immune system sensors might incorrectly identify these dsRNA molecules as foreign, triggering inappropriate IFN responses and hindering cell proliferation. To identify the specific RNA sensors responsible for detecting immunostimulatory RNA and subsequently activating inflammatory responses and arresting cell growth in m^6^A-deficient cells, we performed genetic ablations of key cytoplasmic RNA sensors, including RIG-I, MDA5, and MAVS, in METTL3 KO cells to assess which sensor ablation can mitigate immune responses. Our data indicated that MDA5 deletion did not rescue growth defects in METTL3 KO cells. However, RIG-I or MAVS deletion led to a substantial improvement in cellular proliferation in METTL3 KO cells (Figure 2A). Furthermore, the removal of RIG-I or MAVS can significantly attenuate ISG expression, whereas MDA5 deletion had a minimal effect (Figure 2B). Collectively, these observations underscore the pivotal role of the RIG-I/MAVS signaling activation in mediating the suppressed cellular growth and heightened IFN signaling observed in METTL3 KO cells.

**Figure 2.**
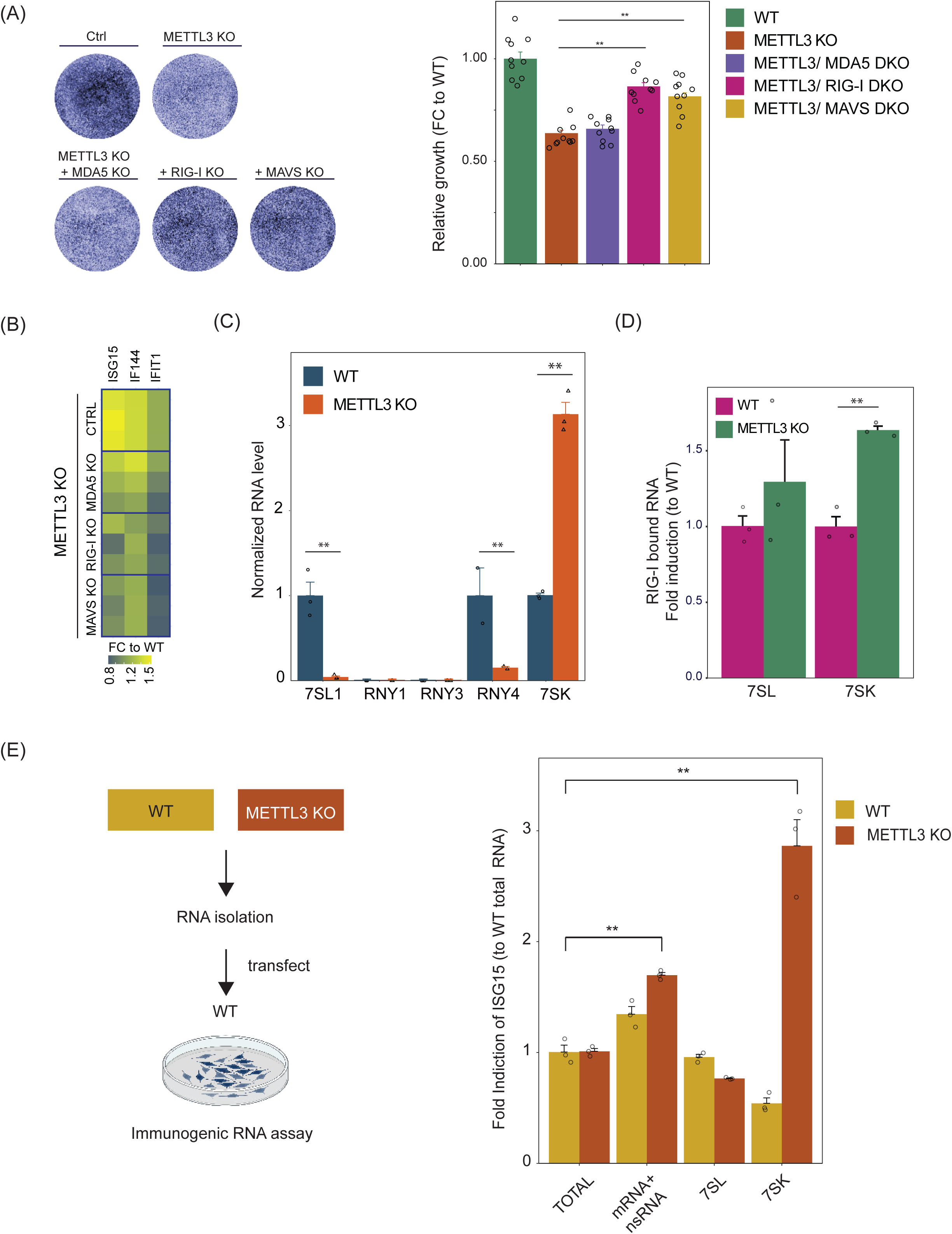
m^6^A deficiency activates the RIG-I/MAVS pathway to elicit inflammatory responses. (A) Crystal violet assays (left) and statistical analysis (right) of growth effects in METTL3 KO cells following ablation of RIG-I, MDA5, and MAVS. Data are normalized to WT and presented as mean ± SEM (n = 5 experimental replicates; **P < 0.01, t-test). (B) Reduction in inflammatory response in METTL3 KO cells upon suppression of RIG-I or MAVS. RT-qPCR results depict changes in the expression of selected ISGs between WT, METTL3 KO, and their combined KO conditions. ISG expression fold changes relative to WT are shown (n = 3 experimental replicates). (C) Comparative analysis of 5′ppp RNA levels between WT and METTL3 KO cells, based on RNA-seq data. Transcripts per million (TPM) values were computed and compared to WT conditions. Results represent average fold changes ± SEM (n = 3 experimental replicates; **P < 0.01, t-test). (D) RIP-qPCR illustrating the differential binding of RIG-I to 7SL and 7SK transcripts in WT and METTL3 KO cells. Data are normalized to WT and expressed as mean ± SEM (n = 3 experimental replicates; *P < 0.01, t-test). (E) Evaluation of the immunostimulatory potential of RNA isolated from METTL3 KO cells compared to WT cells. The schematic on the left outlines the experimental design. ISG15 induction, measured by RT-qPCR, serves as an indicator of the RNA ability to provoke inflammatory responses. Results are normalized to the WT total RNA stimulus and presented as average fold changes ± SEM (n = 3 experimental replicates; *P < 0.05, t-test).

RIG-I is known to recognize and be activated by dsRNA that features a 5’-triphosphate (5’ppp) terminus(21). Early studies have shown that a significant source of endogenous 5’ppp RNA is RNA polymerase III (POL3)-mediated transcription, with increased POL3 activity leading to an overproduction of 5’ppp RNA which then activates RIG-I(22–24). POL3 is responsible for transcribing various noncoding RNAs, including U6 small nuclear RNAs (snRNA), 5S ribosomal RNA, 7SK, 7SL, and Y RNA(23). Despite similar POL3 expression levels between WT and METTL3 KO cells (data not shown) in our RNA-seq data, a detailed examination of POL3-regulated transcripts revealed a significant increase in 7SK expression in METTL3 KO cells, whereas 7SL and YRNAs exhibited decreased expression (Figure 2C). Given previous research suggesting that 7SK can activate RIG-I, we conducted anti-RIG-I RNA immunoprecipitation followed by quantitative PCR (RIP-qPCR) to determine if 7SK exhibited increased interaction with RIG-I in METTL3 KO cells. Our results showed that 7SK has an enhanced binding to RIG-I in the absence of m^6^A compared to WT cells (Figure 2D). Considering that 7SK is extensively methylated with m^6^A(25,26), our finding suggests m^6^A may suppress the interactions between 7SK and RIG-I, thereby inhibiting 7SK immunostimulatory potentials.

To further investigate whether 7SK gains immunostimulatory competence in the absence of m^6^A, thus erroneously activating the immune response, we segregated RNA into total RNA, messenger RNA (mRNA)/non-coding RNA (ncRNA), 7SL, and 7SK from both WT and KO cells. We then introduced identical amounts of RNA (500 ng) into WT cells to evaluate their potential to induce ISG expression. Our results showed that while mRNA/ncRNA from m^6^A-deficient cells could provoke inflammatory responses in WT cells, the effect was comparatively mild. In contrast, 7SK RNA isolated from m^6^A-deficient cells markedly stimulated ISG expression in METTL3 KO cells (Figure 2E). Collectively, our findings indicate that the absence of m^6^A modifications significantly enhances the immunostimulatory capacity of 7SK RNA, subsequently activating the RIG-I/MAVS signaling pathway.

### YTHDC1 modulates 7SK levels and suppresses RNA immunogenicity

m^6^A modifications recruit YTH family proteins to regulate various aspects of RNA metabolism including gene expression, RNA stability, and translation. To determine which YTH family proteins modulate 7SK RNA and influence its immunogenicity, we first analyzed the binding preferences of YTHDF1, YTHDF2, and YTHDC1 to 7SK. This analysis was conducted using recently published reverse transcription-based RBP (RNA-binding protein) binding site sequencing assay (ARTR-seq) datasets in HeLa cells(27). Our analysis revealed that YTHDC1 exhibits significant binding to 7SK RNA (Figure 3A), in contrast to the negligible or absent binding observed for YTHDF1 and YTHDF2.

**Figure 3.**
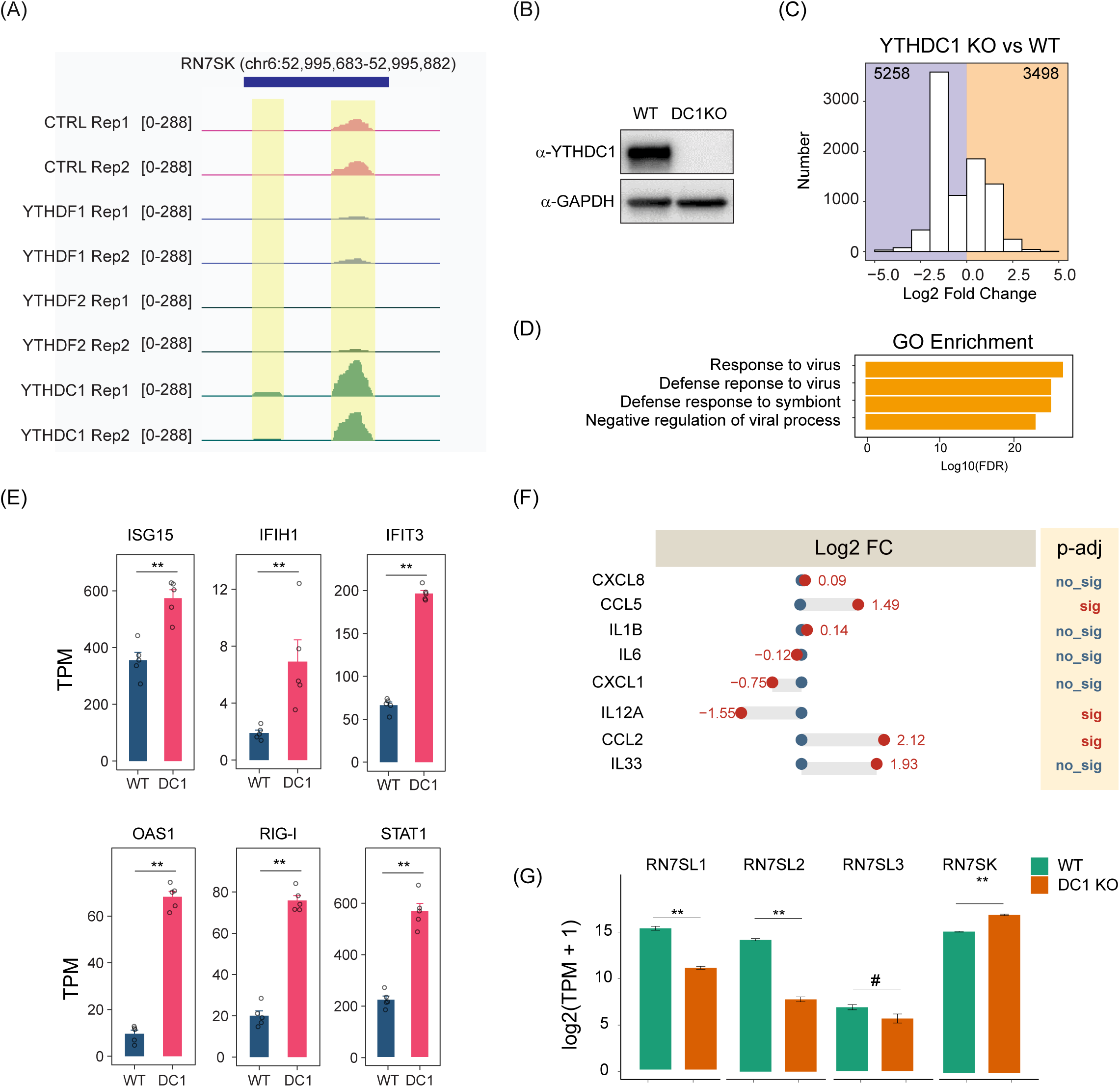
YTHDC1 modulates 7SK levels and suppresses RNA immunogenicity. (A) Snapshots from the Integrative Genomics Viewer (IGV) demonstrating overlaps of YTHDC1 ARTR-seq signal with RN7SK in HeLa cells. (B) Verification of YTHDC1 KO using Western blot analysis. (C) RNA-seq analysis reveals genes differentially expressed between YTHDC1 KO and WT cells. (D) GO analysis illustrating the enrichment of pathways related to antiviral responses among upregulated genes in YTHDC1 KO cells. (E) Expression levels of ISGs in YTHDC1 KO cells, as identified by RNA-seq. Expression values are presented as TPM based on data from three experimental replicates. (F) Mild induction of cytokines and chemokines observed in YTHDC1 KO cells, as determined by RNA-seq. Expression levels are normalized to WT and presented as log2 fold changes. (G) Expression levels of 7SL and 7SK in YTHDC1 KO cells, as identified by RNA-seq. Expression values are normalized to WT and presented as log2 (TPM+1).

To further investigate whether YTHDC1 influences 7SK RNA to modulate innate immune responses, we generated YTHDC1 KO lines in HMC3 cells and assessed the inflammatory responses elicited by YTHDC1 deficiency. Western blot analysis confirmed the absence of YTHDC1 expression in these KO cell lines (Figure 3B). Subsequent RNA-seq analysis revealed significant alterations in gene expression, with 3,498 genes upregulated and 5,258 genes downregulated (Figure 3C). GO analysis of the upregulated genes showed enrichment in innate antiviral immune responses (Figure 3D), accompanied by the induction of ISGs and a broad array of cytokine and chemokine activation (Figure 3E and F), mirroring the gene expression responses observed in m^6^A-deficient cells (Figure 1D-G) (Supplementary Figure S2). Moreover, similar to observations in m^6^A deficient cells, YTHDC1 KO led to an induction of 7SK RNA (Figure 3G). Collectively, our findings suggest that the m^6^A-YTHDC1 axis serves as a crucial regulatory pathway for controlling 7SK RNA levels and its associated immunogenicity, playing a key role in restricting self RNA-activated inflammatory responses.

### m^6^A deficiency promotes ZBP1 oligomerization to induce cell death

m^6^A dysregulation can impact cell growth, differentiation, metabolic activity, and overall cellular function. While various m^6^A-related mechanisms, particularly those impacting RNA stability and protein synthesis, are thought to disrupt key biological regulators, potentially leading to progressive tissue malformation or degeneration(10,28,29), our RNA-seq analysis indicates that elevated inflammatory responses are frequently associated with tissue degeneration in m^6^A KO mice (Supplementary Figures S3A). Furthermore, a significant increase in ZBP1, which is involved in both necroptotic and apoptotic cell death pathways, was consistently observed in tissues from m^6^A KO mice exhibiting heightened IFN responses and increased cell death (Supplementary Figures S3A and B). These findings led us to hypothesize that prolonged inflammation, in conjunction with ZBP1 activity, might intensify cell death in m^6^A-deficient cells (Figure 4A).

**Figure 4.**
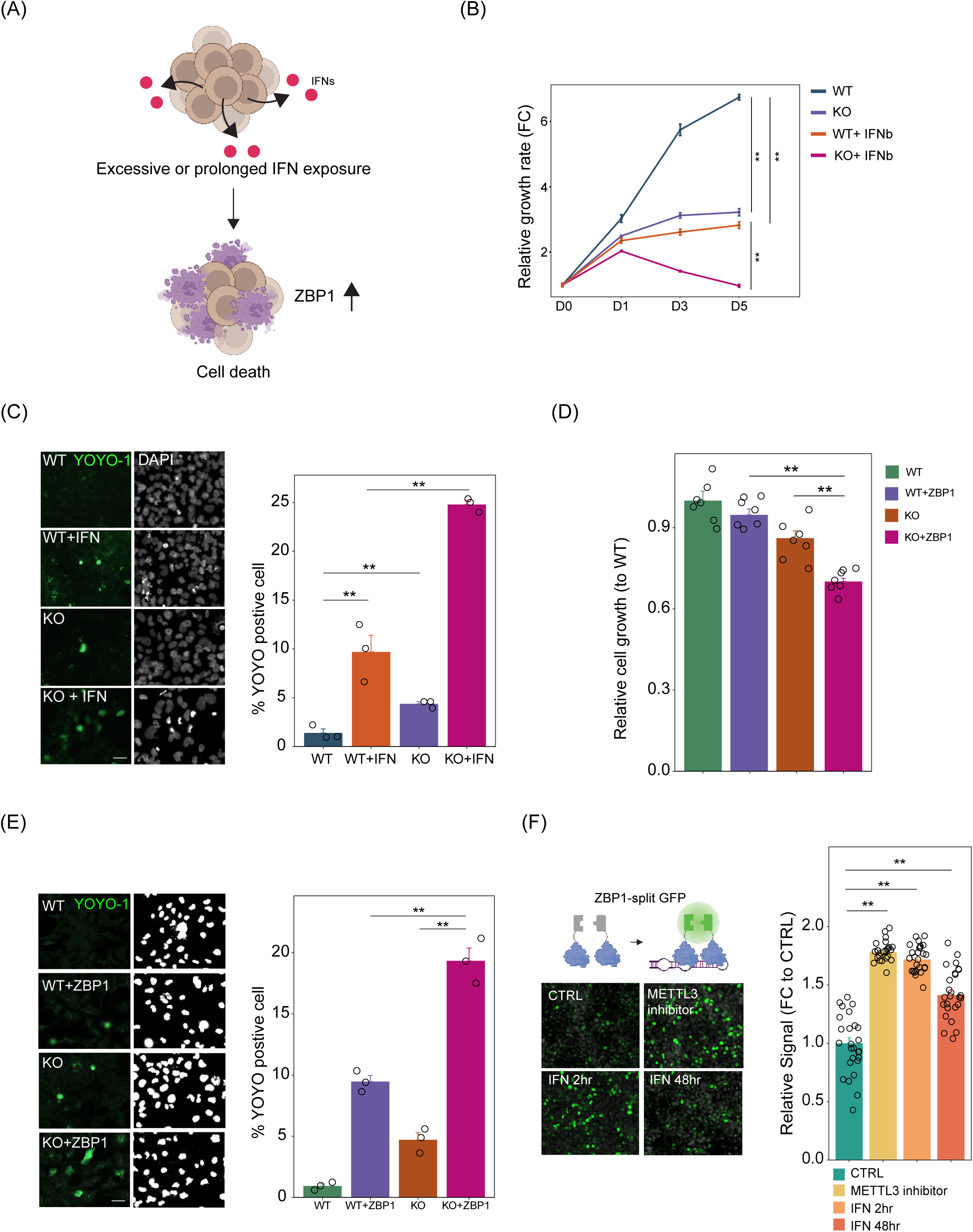
m^6^A deficiency promotes ZBP1 oligomerization to induce cell death. (A) Illustrative model showing how excessive or prolonged IFN exposure exacerbates cell death in m^6^A-deficient cells. (B) Analysis of cell proliferation in WT and METTL3 KO cells treated with either a vehicle or IFN. Growth values are normalized to WT at Day 0 (D0) and displayed as mean fold change ± SEM (n = 5 experimental replicates; **P < 0.01; two-way ANOVA). (C) YOYO-1 staining used to quantify dying cells in WT and METTL3 KO cells, with and without IFN treatment. Results are presented as mean ± SEM (n = 3 experimental replicates; **P < 0.01; t-test). (D) Comparison of cell growth in WT and METTL3 KO cells upon ZBP1 overexpression. Data are normalized to WT and shown as mean fold change ± SEM (n = 5 experimental replicates; **P < 0.01; t-test). (E) Representative images and quantitative analysis of cell death in WT and METTL3 KO cells following ZBP1 overexpression. Values are presented as mean ± SEM (n = 3 experimental replicates; **P < 0.01; t-test). (F) Assessment of ZBP1 oligomerization under various treatment conditions using the ZBP1-BiFC assay. Results are normalized to control BiFC intensity and displayed as mean fold change ± SEM (n = 25 cells per condition; **P < 0.01; t-test).

To test this hypothesis, we exposed both WT and METTL3 KO cells to IFNs and monitored how elevated and prolonged inflammation affects their growth over five days. The METTL3 KO cells exhibited a substantial decrease in growth compared to WT cells, and this growth inhibition became even more pronounced following IFN treatment after 3 days, suggesting that prolonged inflammation could exacerbate cell death in METTL3 KO cells (Figure 4B). To further validate these findings, we used YOYO-1 staining to identify apoptotic cells and found an increased number of YOYO-1 positive cells in the METTL3 KO condition after 48 hours of IFN exposure (Figure 4C). Together, these results confirm that m^6^A-deficient cells are more susceptible to cell death under conditions of excessive and prolonged inflammation.

Our RT-qPCR analysis confirmed that IFN exposure significantly induced ZBP1 expression in METTL3 KO cells (Supplementary Figure S3C), aligning with the observed elevation of ZBP1 in cell death scenarios in m^6^A KO animals. To further examine the role of ZBP1 in promoting cell death under m^6^A deficiency, we overexpressed ZBP1 in both WT and METTL3 KO cells and then conducted growth assays and YOYO-1 staining to assess cell viability. Interestingly, while ZBP1 overexpression led to cell death in WT cells, mirroring the effects seen with IFN treatment (Figures 4D and E), METTL3 KO cells showed an even more pronounced increase in cell death following ZBP1 overexpression. These findings lead us to hypothesize that m^6^A deficiency may enhance the formation of ZBP1 RNA ligands, intensifying ZBP1 activation and subsequently contributing to increased cell death in m^6^A KO cells.

ZBP1 can dimerize or oligomerize upon binding to its nucleic acid substrates, which then initiates a signaling complex capable of inducing inflammation or cell death(30,31). We utilized a ZBP1 bimolecular fluorescence complementation (BiFC) assay to further explore whether m^6^A deficiency increased the levels of ZBP1 RNA substrates, thereby enhancing ZBP1 oligomerization (Figure 4F). This assay was conducted in 293T cells, which lack an active downstream ZBP1 signaling pathway, to avoid confounding effects from cell death. Our findings revealed that treatment with the METTL3 inhibitor STM2457(32), which blocks m^6^A RNA methylation, led to an increased BiFC signal (Figure 4F). Additionally, short-term IFN treatment also elevated the BiFC signal. Collectively, our results suggested both m^6^A deficiency and IFN exposure contribute to the production of ZBP1 RNA ligands, thereby promoting ZBP1 oligomerization and leading to cell death.

### Z**α** domains interact with transcripts harboring RNA G-quadruplex (rG4)

We next sought to identify specific RNA ligands of ZBP1 to better understand how ZBP1 potentially facilitates cell death under m^6^A deficiency conditions. ZBP1 harbors Zα domain that recognize Z-RNA to elicit downstream cell death pathways(33). Identifying these ZBP1 RNA ligands poses a significant challenge, as the cell death induced by ZBP1 expression complicates the precise determination of its RNA targets using conventional CLIP-seq. To circumvent this, we strategically utilized the longer isoform of ADAR1, ADAR1 P150, which, alongside ZBP1, is the only known protein in the human proteome with Zα domain(34), to identify potential Z-RNA (Figure 5A).

**Figure 5.**
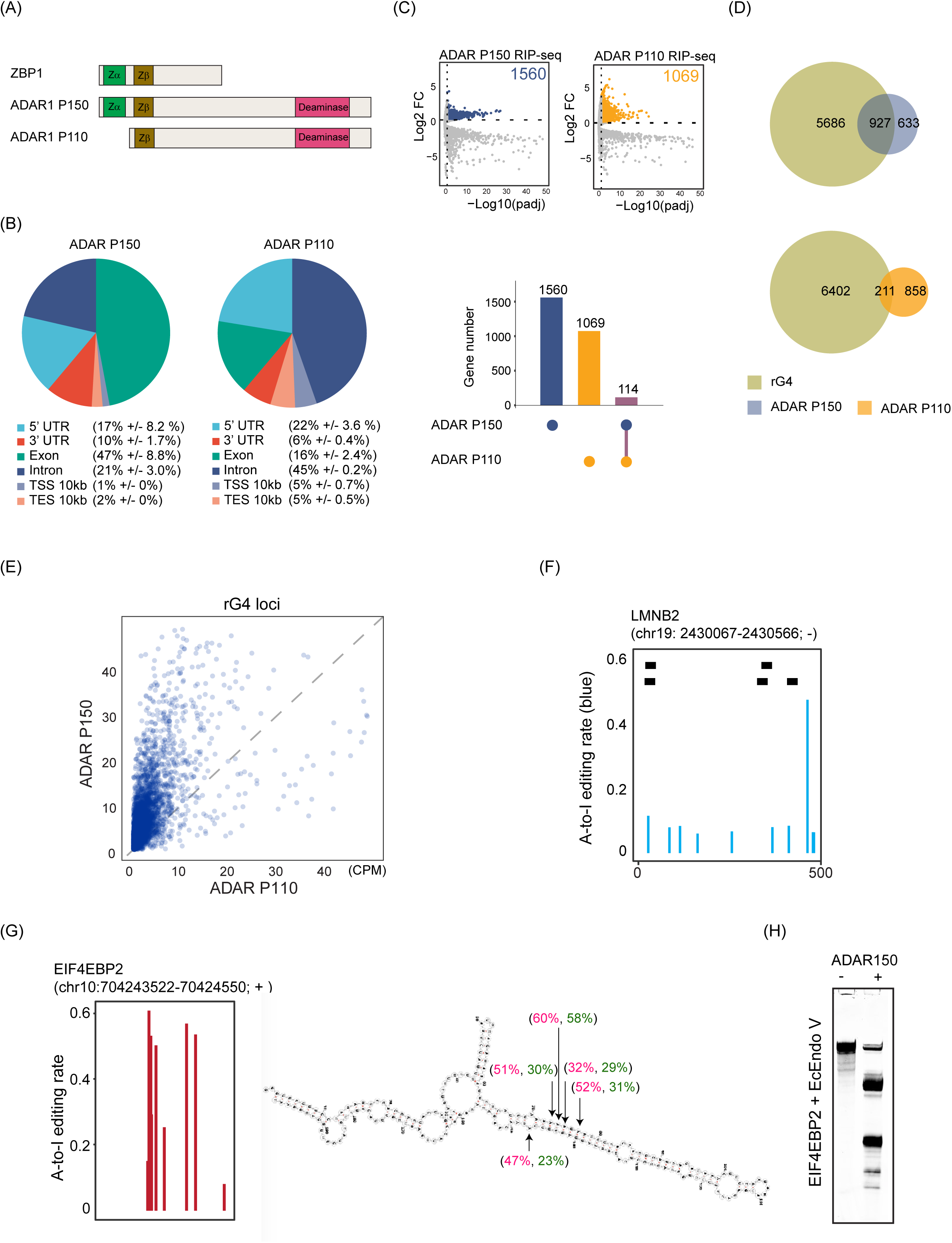
Zα domains interact with transcripts harboring RNA G-quadruplex (rG4) Structures. (A) Illustrative schematic depicting the protein domain structures of ZBP1, ADAR1 P150, and P110. (B) RIP-seq analysis of ADAR1 P110 and ADAR1 P150, showing distinct distributions of sequence reads across various transcriptomic regions. (C) Comparative analysis identifying specific RNA transcripts bound by ADAR1 P110 and ADAR1 P150. Overlap analysis indicates minimal shared transcripts between the two isoforms. (D) Venn diagram illustrating the correlation between rG4-containing transcripts and those bound by ADAR1 P110 and P150. Notably, ADAR1 P150 shows a significantly higher overlap (about 50%) with rG4-rich transcripts compared to ADAR1 P110 (under 20%). (E) Scatter plot demonstrating that sequence reads from ADAR1 P150 are more prevalent in rG4 regions than those from ADAR1 P110. (F) Visualization of A-to-I editing near rG4 regions in LMN2 transcripts. (G) The 3’UTR of EIF4EBP2 featuring a dense cluster of A-to-I editing. EIF4EBP2 3’UTR, which was predicted to form a duplex RNA structure, harbored extensive A-to-I editing (red: revealed by RNAseq data; green: confirmed by Sanger sequencing). (H) Validation of the EIF4EBP2 3’UTR as a substrate of the Zαb domain protein. The EIF4EBP2 3’UTR was confirmed as a substrate for ADAR1 P150 using an in vitro Endo V enzymatic assay.

Specifically, ADAR1 P150 has the unique capability to competitively bind with ZBP1 RNA ligands, thus mitigating ZBP1-induced cell death(35,36). This attribute makes ADAR1 P150 an invaluable tool for exploring the RNAs that interact with ZBP1, helping to reveal potential mechanisms of action. In contrast, the shorter isoform, ADAR1 P110, which lacks the Zα domain, served as a negative control in our experiments.

To this end, we analyzed the RIP-seq data for ADAR1 isoforms P110 and P150, which revealed distinct RNA substrates for each (Figures 5B and C). Specifically, sequencing reads for ADAR1 P110 predominantly occurred in intronic regions, while reads from ADAR1 P150 were enriched in exonic and 3’UTR regions (Figure 5B). This distinction is further reflected in their A-to-I editing patterns, where ADAR1 P110 primarily edits introns, and ADAR1 P150 edits are mainly found in exonic and 3’UTR areas (Supplementary Figures S4A and B). Interestingly, the editing activity of ADAR1 P150 is somewhat lower than that of P110 (Supplementary Figure S4C), suggesting that the Zα domains in ADAR1 P150 may guide the protein to specific RNA sites, which may not necessarily be the optimal substrates for its enzymatic action.

Our analysis of the ADAR1 P110 and P150 RIP-seq datasets showed that each isoform binds to a distinct set of enriched transcripts, with minimal overlap between them (Figure 5C and Supplementary Table 2). Given prior research indicating that Zα domains interact with DNA G-quadruplexes (G4) (37), we hypothesized that RNAs with G-quadruplexes (rG4) might similarly serve as substrates for ADAR1 P150 and ZBP1. To explore this hypothesis, we compared the RNAs associated with ADAR1 P150 and P110 to the known human rG4 RNA dataset(38). Our analysis revealed that approximately 60% of transcripts associated with ADAR1 P150 contained rG4 structures, whereas only about 20% of ADAR1 P110-associated transcripts had such structures (Figure 5D and Supplementary Table 2). Additionally, sequence reads from ADAR1 P150 were significantly more prevalent in rG4 regions compared to those from ADAR1 P110 (Figure 5E).

LMNB2 stands out as one of the RNA ligands for Zα domain, where the binding enrichment of ADAR1 P150 was notably apparent. This transcript not only contains rG4 structures but also features several A-to-I editing sites in close proximity to these regions (Figure 5F). Additionally, we observed that some duplexed RNA, such as EIF4EBP2, features a cluster of highly edited A-to-I sites in its 3’UTR and can also serve as an enzymatic substrate for ADAR1 P150 (Figures 5G-H and Supplementary Table 3). Collectively, our findings suggest that Zα-domain proteins can interact with both rG4 RNA and duplexed RNA.

### ZBP1 engages with RNA transcripts that possess G-quadruplex structures

To further confirm our findings, we employed the Surveying Targets by APOBEC Mediated Profiling (STAMP) technique(16) to orthogonally identify RNA targets of ZBP1. We engineered a plasmid to express ZBP1 Zαβ domain with APOBEC1 (hereafter ZBP1zαβ-APOBEC1). ZBP1 Zαβ domain directs APOBEC1 to specific RNA targets, facilitating C-to-U base conversion near the actual ZBP1zαβ RNA binding loci (Figure 6A).

**Figure 6.**
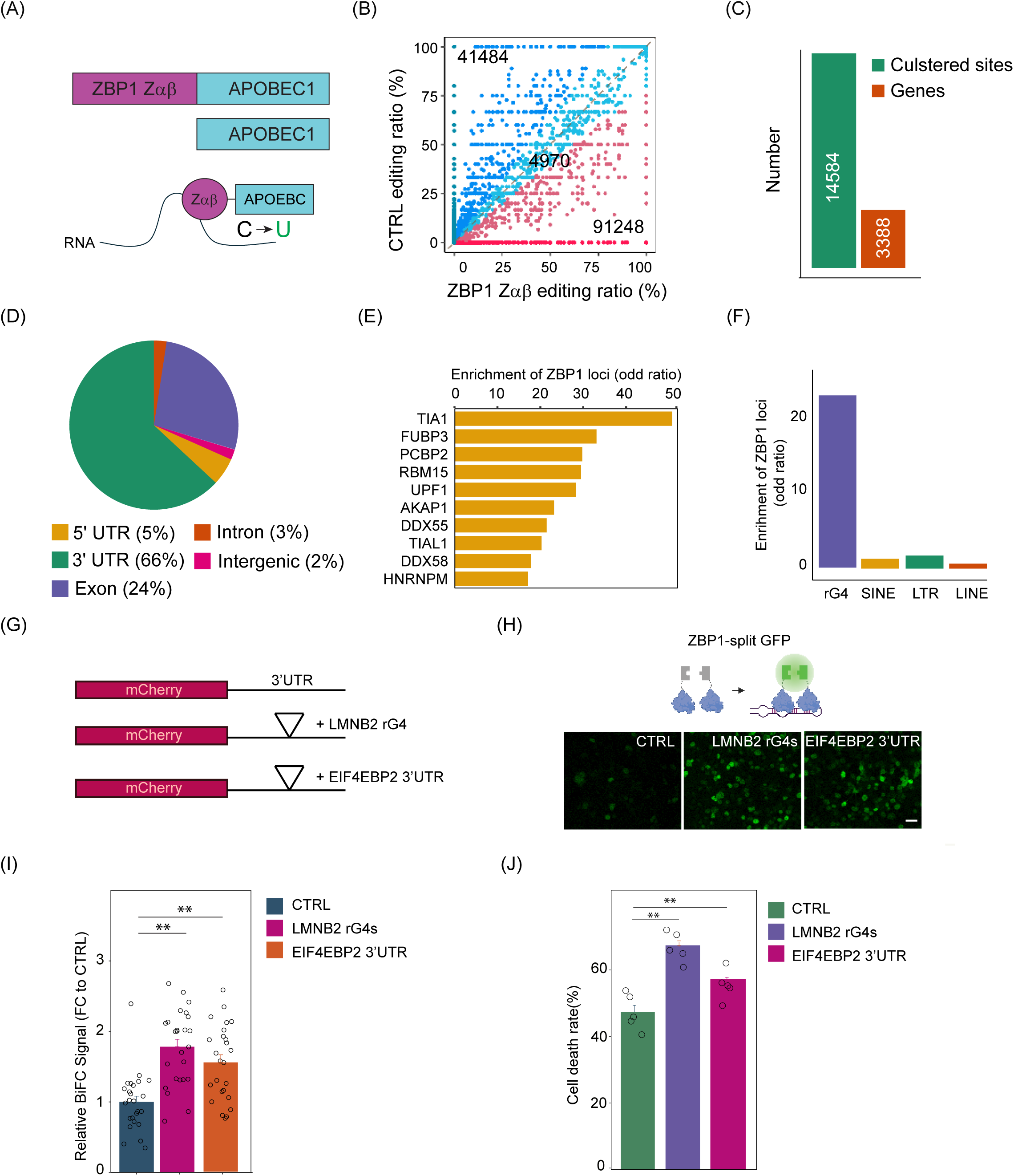
ZBP1 engages with RNA transcripts that possess G-quadruplex structures and induces cell death. (A) Schematic illustration of the ZBP1-STAMP method, where APOBEC1 is fused to ZBP1 to guide C-to-U editing at cytidine residues near ZBP1 RNA binding sites. (B) A scatter plot comparing the distribution of C-to-U editing sites in cells overexpressing ZBP1Zαβ-APOBEC1 versus APOBEC1 alone, highlighting the distinct editing distributions between ZBP1-STAMP and control-STAMP (background). (C) Bar graph displaying the number of clustered C-to-U editing sites and the genes targeted by ZBP1Zαβ--APOBEC1. (D) Pie charts illustrating the distribution preferences of ZBP1-APOBEC1 mediated C-to-U editing across different transcript regions. (E) Enrichment analysis of RNA-binding proteins (RBPs) at ZBP1Zαβ-APOBEC1 mediated C-to-U editing sites, with the odds ratio calculated to determine the enrichment score (**P < 0.01; Fisher exact test). (F) Enrichment analysis of rG4 sites and other repetitive elements at ZBP1-APOBEC1 mediated C-to-U editing sites, with the odds ratio calculated for the enrichment score (**P < 0.01; Fisher exact test). (G) Schematic representation of the plasmid design. The 3’ UTRs of LMNB2 and EIF4EBP2 were incorporated into the mCherry 3’ UTR region of the AAV-mCherry expression vector. (H) Representative images demonstrating increased ZBP1 oligomerization in cells transfected with the 3’ UTRs of LMNB2 and EIF4EBP2 (scale bar: 200 μm). (I) Quantitative analysis of ZBP1-BiFC assay upon expression of the 3’ UTRs of LMNB2 and EIF4EBP2. Results are normalized to the control and presented as mean fold change ± SEM (n = 30 cells per condition across two independent experimental replicates; **P < 0.01; t-test). (J) Quantitative analysis of cell death in ZBP1 overexpressing cells that were subjected to different transfection. Cell death percentages are reported as mean ± SEM (n=5; (**P < 0.01; t-test)

We transiently transfected cells with ZBP1zαβ-APOBEC1 and a control APOBEC1, then subjected them to RNA sequencing 48 hours after transfection. We restricted at least 10x coverage and a minimum of 5% C-to-U editing ratio, to determine confident editing events. After excluding editing sites found in the APOBEC1 control samples, we identified 91,248 editing sites specific for ZBP1zαβ-APOBEC1 (Figure 6B). By clustering editing sites that were within 100 bases of each other, this analysis yielded 14,584 editing clusters, representing 15.9% of the initial unfiltered windows, distributed across 3,388 transcripts (Figure 6C and Supplementary Table 4), with over 50% of these clusters located within the 3’ untranslated regions (3’ UTRs) of the transcripts (Figure 6D).

We further analyzed published RBPs CLIP-seq datasets to determine whether any RBPs co-localize with ZBP1zαβ binding sites that could potentially influence ZBP1 function(39). Our analysis revealed significant enrichment of TIA1, FUBP3, PCBP2, and RBM15 RNA binding sites overlaps with ZBP1zαβ editing clusters (Figure 6E). Supporting our analysis, previous studies have reported that TIA1 can colocalize with ZBP1 in stress granules(40). Our findings suggest that several RBP candidates may regulate how ZBP1 interacts with its RNA targets, and ZBP1 association with RNA in stress granules may play a role in inducing necroptosis.

Lastly, we investigated whether ZBP1zαβ interacts with rG4 RNA and ERV elements, which had been previously proposed as Z-RNA(41–43). Unexpectedly, our analysis revealed minimal overlap between ZBP1zαβ editing clusters and ERV elements (Figure 6F). In contrast, we found significant enrichment of rG4 RNA within ZBP1zαβ editing clusters. Together, our findings from the ADAR P150 RIP-seq datasets and ZBP1zαβ editing profiles suggest that rG4 RNAs are potential ligands for Zαβ domain-containing proteins.

### rG4 RNA promotes ZBP1 oligomerization and cell death

To further validate the functional interaction between rG4 RNA and ZBP1, we incorporated the 3’ UTRs of LMNB2, containing rG4 structures, into the mCherry 3’UTR of the AAV-mCherry expression vector (Figure 6G). Subsequently, we assessed whether overexpression of this construct could induce ZBP1 oligomerization using the ZBP1 BiFC assay. Remarkably, the LMNB2 rG4s significantly enhanced ZBP1 BiFC signals compared to the control mCherry 3’UTR. Intriguingly, the 3’ UTR of EIF4EBP2, identified as a substrate for ADAR150 and ZBP1 in our analysis and predicted to form RNA duplexes, also effectively promoted ZBP1 oligomerization (Figure 6H and 6I).

To explore whether rG4 and duplexed RNA can activate ZBP1-mediated cell death, we co-transfected these constructs along with ZBP1-expressing plasmids into HeLa cells. Subsequent cytotoxicity assays showed that both rG4 and duplexed RNA significantly increased cell death compared to the control mCherry 3’UTR (Figure 6J). Collectively, these results suggest that both rG4 and duplexed RNA can serve as ZBP1 RNA ligands to induce cell death.

### Deficiency in the m^6^A-YTHDF1/2 axis promotes the rG4 formation during in IFN exposure

As our findings suggest ZBP1 interacts with rG4-containing transcripts, we hypothesized that conditions such as m^6^A deficiency or IFN exposure might promote widespread rG4 formation in the transcriptome, thereby enhancing ZBP1 to bind and mediate cell death. To test this hypothesis, we used a hemin-based biotin assay where rG4 interacts with hemin to catalyze biotin-phenol, producing red fluorescence indicative of rG4 levels(19), allowing us to quantify rG4 in METTL3 KO and IFN-exposed conditions (Figure 7A). Our data showed that METTL3 KO and IFN treatment both promote rG4 formation (Figure 7B and 7C). Furthermore, the combination of METTL3 KO and IFN treatment significantly increased rG4 production, correlating with our observations of heightened cell death when these two conditions are combined (Figure 7B and 7C). Taken together, these observations support the notion that METTL3 KO encourages rG4 formation that favor ZBP1 binding and ultimately leading to the activation of cell death pathway.

**Figure 7.**
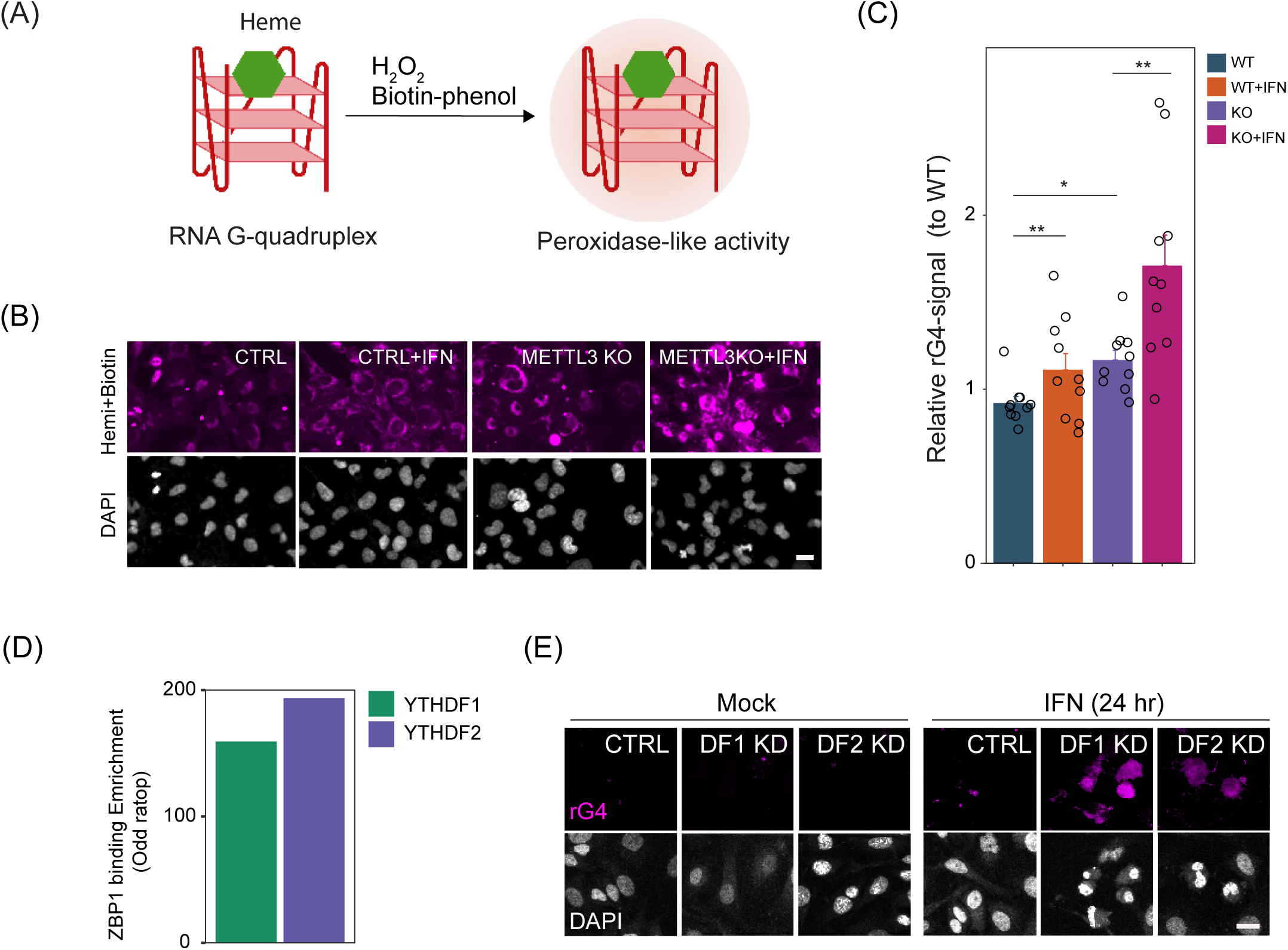
Deficiency of m^6^A-YTHDF1/2 axis encourages rG4 formation in the presence of IFN exposure. (A) Illustration of the hemin-based biotin assay used to detect rG4 formation. In this assay, rG4 structures interact with hemin, catalyzing the reaction with biotin-phenol and producing red fluorescence to indicate the presence of rG4. (B) Representative images showing increased rG4 formation in cells subjected to METTL3 KO, IFN treatment, or both (scale bar: 10 μm). (C) Quantitative analysis demonstrating enhanced rG4 formation following METTL3 KO, IFN treatment, or their combination. Results are normalized to the control and presented as mean fold change ± SEM (n = 10 cells per condition across two independent experimental replicates; **P < 0.01; t-test). (D) Enrichment analysis showing the association of YTHDF1 and YTHDF2 RNA occupancy with ZBP1 binding sites (**P < 0.01; Fisher exact test). (E) Hemin-based biotin assay illustrating increased rG4 formation in cells with YTHDF1 or YTHDF2 knockdown (KD) under IFN treatment (n = 3 experimental replicates; scale bar: 20 μm).

To further explore how m^6^A deficiency leads to the accumulation of rG4 RNA, we hypothesized that m^6^A might recruit m^6^A-binding proteins, such as those from the YTH family, to inhibit rG4 formation, thereby limiting ZBP1 to access to its RNA ligands. Supporting this hypothesis, our analysis of existing PAR-seq datasets(17,18) showed that ZBP1 edit clusters are enriched at YTHDF1 and YTHDF2 binding loci (Figure 7D), with 13% of ZBP1 edit clusters containing YTHDF1 binding sites and 15% overlapping with YTHDF2 binding sites (data not shown). This suggests that YTHDF1 and YTHDF2 may play roles in modulating the availability of RNA substrates to ZBP1 by influencing rG4 structure formation.

To further validate this mechanism, we conducted siRNA-mediated knockdown (KD) of YTHDF1 and YTHDF2, followed by either mock or IFN treatment for 24 hours. We then used a hemin assay to assess rG4 formation. Remarkably, similar to m^6^A-deficient cells, KD of YTHDF1 and YTHDF2 alone slightly increased rG4 formation; however, IFN treatment significantly amplified rG4 formation in these KD cells (Figure 7E). Collectively, these findings suggest that the m^6^A-YTHDF1/YTHDF2 axis acts as a protective barrier that prevents transcripts from folding into rG4 structures upon IFN exposure. In the absence of m^6^A or YTHDF1/YTHDF2, ZBP1 can more readily access and bind these regions, potentially leading to its dimerization and subsequent activation. These results highlight a novel regulatory role for the m^6^A-YTHDF1/YTHDF2 axis in modulating ZBP1 activation and suggest a new mechanism to prevent self-nucleic acids from activating inflammatory pathways.

## Discussion

While synthetic RNA inherently exhibits immunogenic properties, the cooperation of modified nucleosides, including m^6^A, m5C, and pseudouridine, allows these synthetic RNAs to evade immune detection and mitigate inflammatory responses within host cells(44). These modifications, also found naturally in the eukaryotic transcriptome, suggest a role in diminishing the immunogenic potential of host RNA, thereby preventing autoimmune responses.

In our studies, we have demonstrated that m^6^A methylation is crucial in suppressing the activity of host immunostimulatory RNA. The absence of m^6^A triggers IFN responses, increasing susceptibility to inflammatory cell death. Additionally, the impact of m^6^A absence in promoting inflammatory responses is evident across various cell types and tissues, as highlighted by our observations of increased ISG expression in diverse tissues from METTL3 KO mice (Supplementary Figure S3A) and in different human cell lines (data not shown). It is important to note, however, that the onset and magnitude of these inflammatory responses can vary, depending on the efficiency and integrity of the innate immune signaling pathways of the cells.

While deficiencies in A-to-I editing are known to activate MDA5-mediated antiviral immune responses, our study indicates that the absence of m^6^A predominantly triggers the RIG-I pathway, leading to inflammatory reactions. Given that MDA5 recognizes long dsRNAs and RIG-I preferentially interacts with short dsRNAs, our findings imply that A-to-I and m^6^A modifications selectively attenuate the immunogenic potential of different dsRNA categories in transcriptome to suppress host RNA immunogenicity. Supporting this, we observed that 7SK acquires immunostimulatory properties following m^6^A depletion, unlike dsRNA derived from Alu repeat elements, which become immunogenic due to a lack of A-to-I editing. Notably, knocking out MDA5 in m^6^A-deficient cells partially rectifies growth abnormalities and reduces the induction of ISGs, suggesting that a subset of long dsRNAs might become immunogenic in the absence of m^6^A to activate MDA5 signaling pathways. Recent studies have identified m^6^A modifications in long interspersed elements (LINEs), short interspersed elements (SINEs), and endogenous retroviruses (ERVs) (20,45). Further investigation is needed to determine whether A-to-I and m^6^A modifications collaboratively modulate the immunogenic characteristics of these RNA types.

m^6^A deficiency has been linked to developmental abnormalities and cell death, often attributed to disruptions in RNA stability and alterations in the translation of critical genes. Our study introduces a unique perspective, suggesting that m^6^A deficiency not only elevates IFN responses but also promotes ZBP1-mediated inflammatory cell death (Figure 8). Mechanistically, our data indicate that m^6^A depletion intensifies inflammation and increases the presence of rG4, which enhances ZBP1 binding and subsequent cell death. This deficiency may reduce the restraints over transcripts adopting rG4 configurations, potentially due to insufficient recruitment of m^6^A-binding proteins such as YTHDF1 and YTHDF2. Moreover, given the observed decrease in m^6^A levels and heightened IFN responses in neurodegenerative diseases, our findings provide potential mechanistic insights into how m^6^A deficiency contributes to the pathogenesis of neurodegeneration. Further exploration of this pathway in neurogenerative disease models, particularly focusing on whether inhibiting the formation of these problematic RNA species or blocking ZBP1 function could open new avenues for mitigating disease progression.

**Figure 8.**
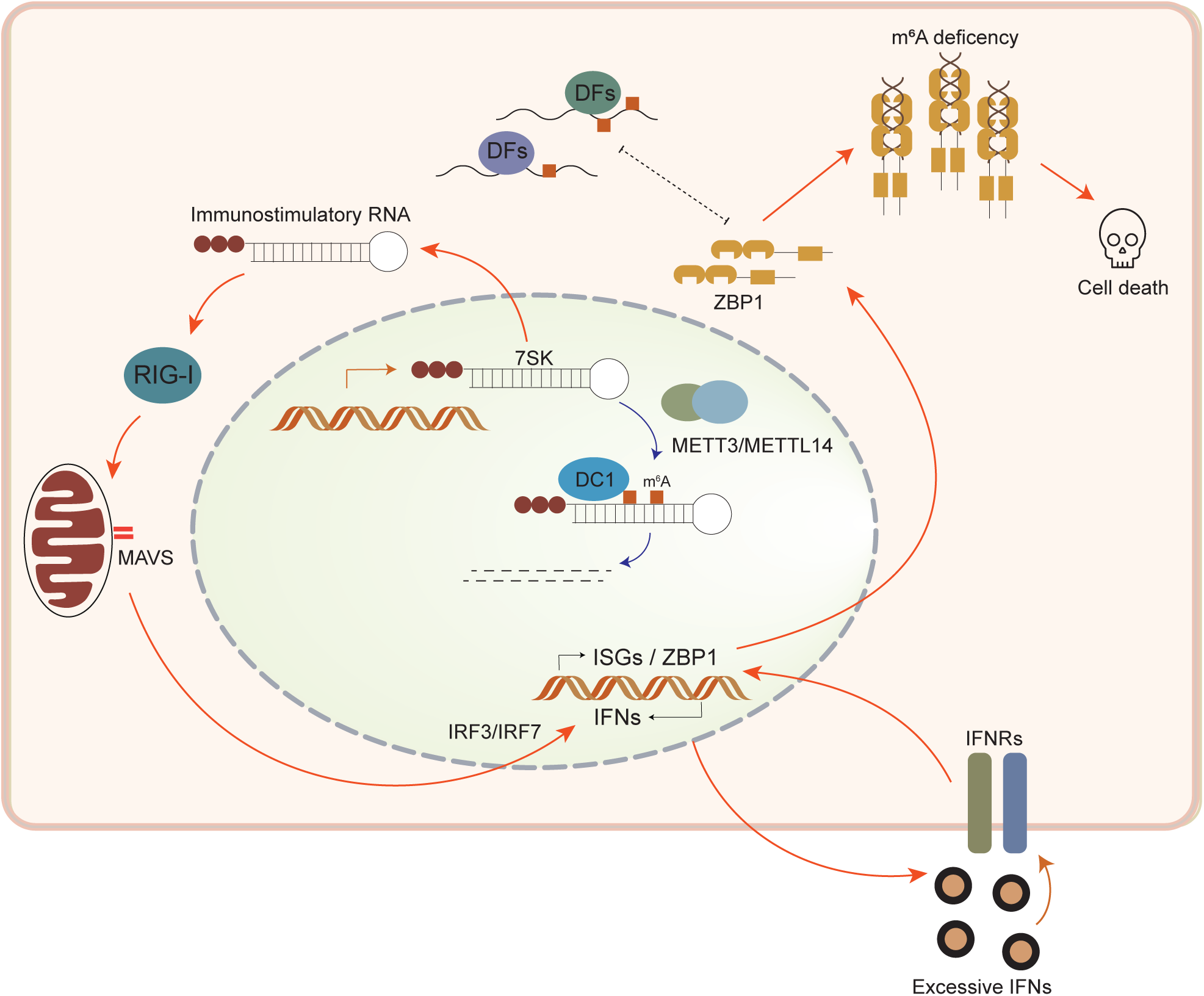
Proposed model for m6A deficiency mediated inflammatory responses and cell death. The scheme illustrating the molecular mechanism by which deficiency in m6A RNA methylation promotes ZBP1-mediated cell death.

## Data availability

RNA-seq data have been deposited to the GEO database (GSE254314) and are publicly available as of the date of publication. Accession numbers are listed in the Supplementary Table 5. This paper does not report the original code. Any additional information required to reanalyze the data reported in this paper is available from the lead contact upon request.

## Acknowledgements

We thank the sequencing support from the Cancer Prevention and Research Institute of Texas (CPRIT RP180734). We express our gratitude to the members of the Weng lab for their critical review of this manuscript and to Anna Dodson for her editorial contributions.

## Author contributions

S.L. and Y-L. W. designed the project and wrote the manuscript. S.L., R.B., constructed libraries for RNA sequencing. Y.C., J-R L., L.S, performed RNA sequencing and A-to-I editing analysis. S.L., X.D, D.P., Y.C., R.B., performed experiments and collected data. G.W.B., Y-L.W. provided administrative support. S.L., Y-L. W. All authors helped edit and contribute to the final version of the manuscript.

---

**Figure S1.**
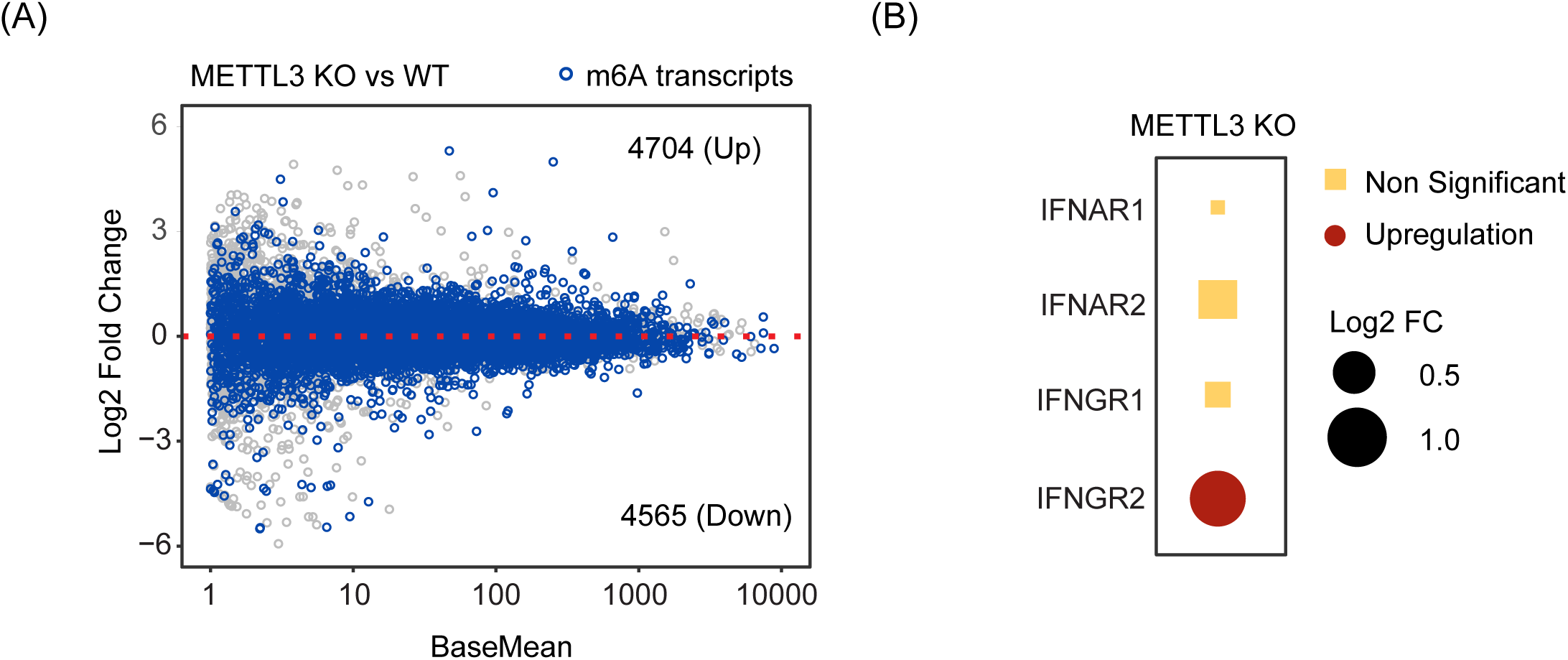
ISG Induction in METTL3 KO cells is not attributed to enhanced RNA stability or IFN receptor upregulation. (A) Volcano plot illustrating differential gene expression between METTL3 KO and WT cells. Transcripts known to undergo m^6^A modifications are highlighted in blue, showing that METTL3 KO does not predominantly result in the upregulation of m^6^A-modified transcripts. (B) Analysis of interferon receptor expression levels in METTL3 KO cells. Expression values are normalized to WT and presented as log2 fold change.

**Figure S2.**
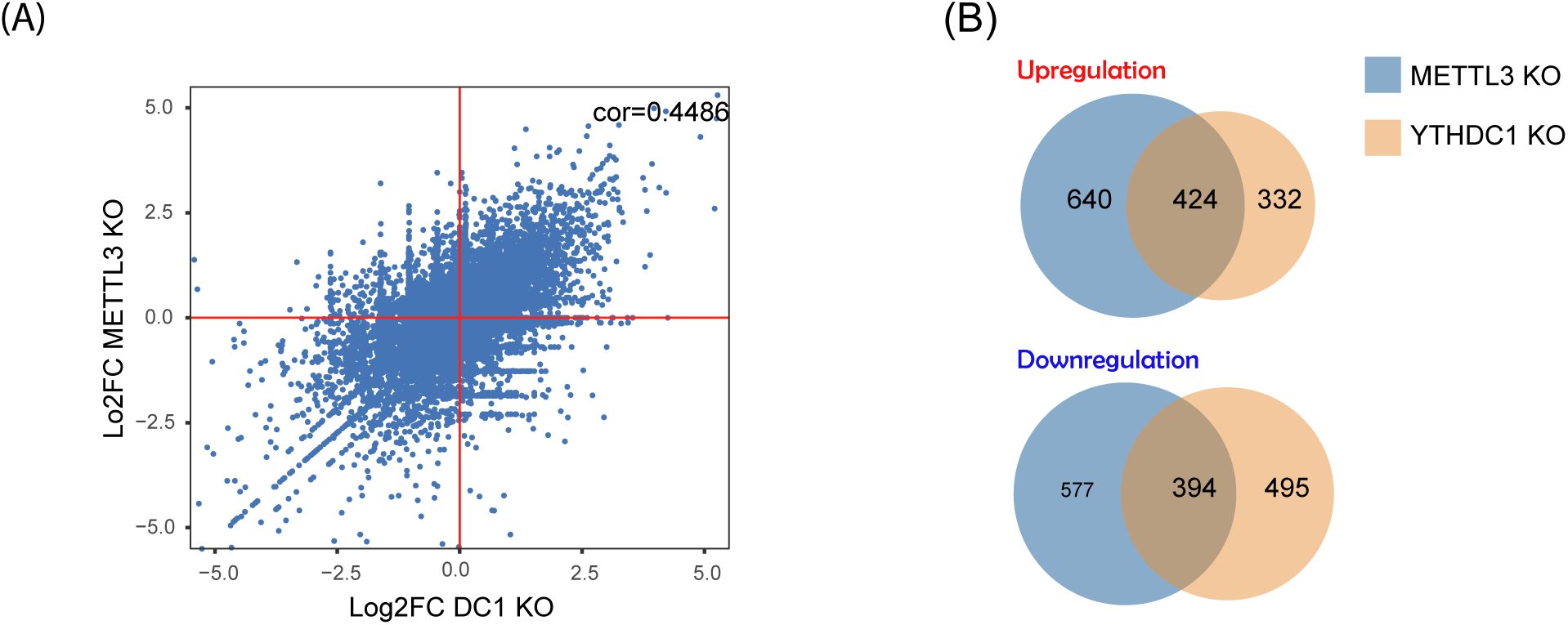
METTL3 and YTHDC1 KOs exhibit similar gene expression changes. (A) Scatter plot illustrating a positive correlation between gene expression changes in METTL3 KO and YTHDC1 KO cells compared to WT cells (r = 0.4486; Pearson’s correlation test). (B) Venn diagram displaying the overlap of genes that are similarly upregulated and downregulated in METTL3 KO and YTHDC1 KO cells relative to WT cells.

**Figure S3.**
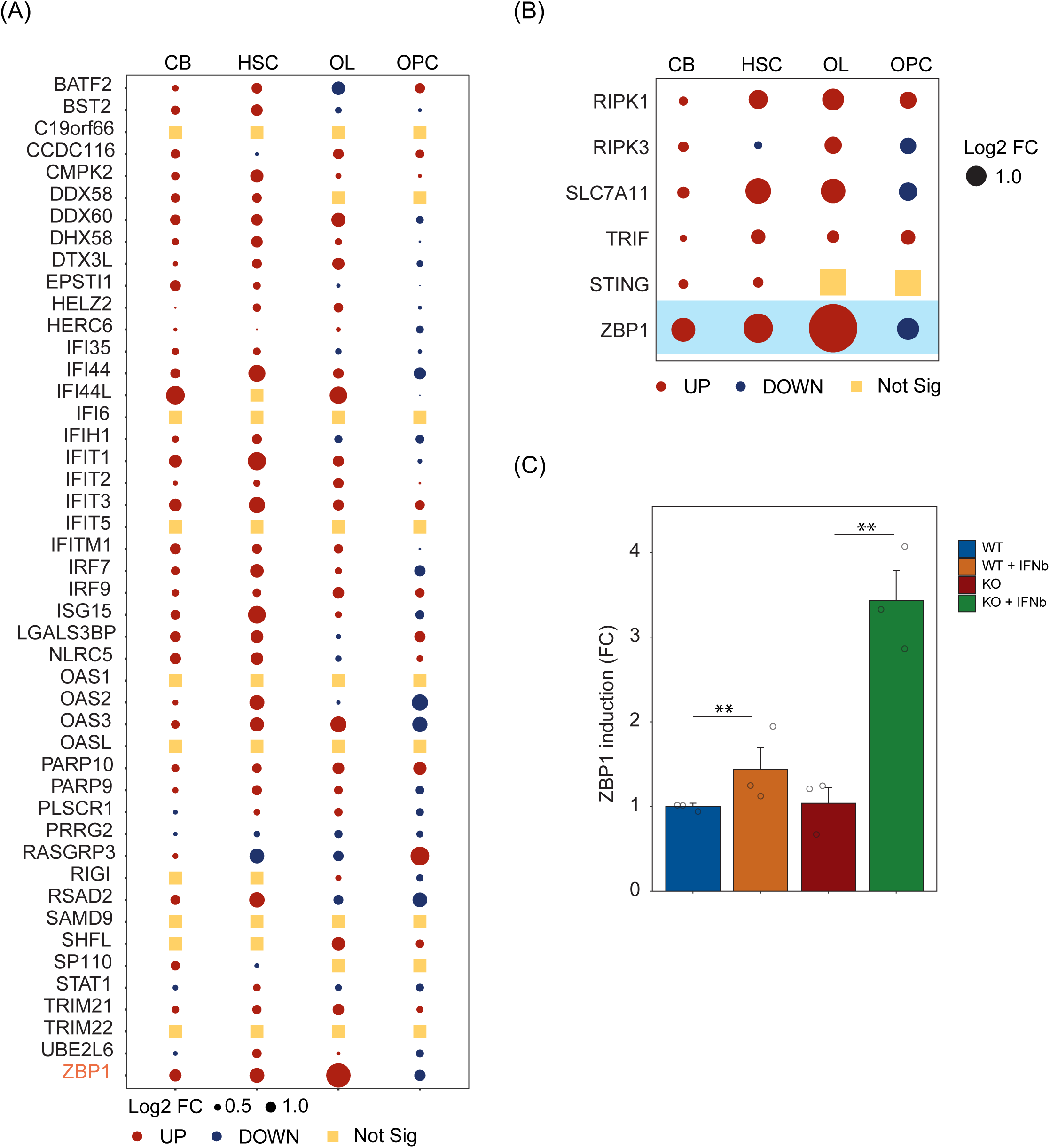
m^6^A deficiency in mice triggers inflammatory responses. (A) Transcriptomic analysis from RNA sequencing (RNA-seq) indicating a marked upregulation of interferon-stimulated genes (ISGs) across various degenerating tissues in m^6^A deficient mice. The affected tissues include the cerebellum (CBs), hematopoietic stem cells (HSCs), and oligodendrocytes (OL), in contrast to non-degenerating oligodendrocyte progenitor cells (OPCs). (B) Dot plot representation illustrating elevated ZBP1 expression levels in degenerating tissues from m^6^A KO mice, highlighting the association of ZBP1 with inflammatory responses in these specific tissues. (C) RT-qPCR analysis measuring ZBP1 expression levels in both WT and METTL3 KO cells, before and after IFN treatment. Data presented as mean ± SEM (n = 3 experimental replications; **P < 0.01, t-test)

**Figure S4.**
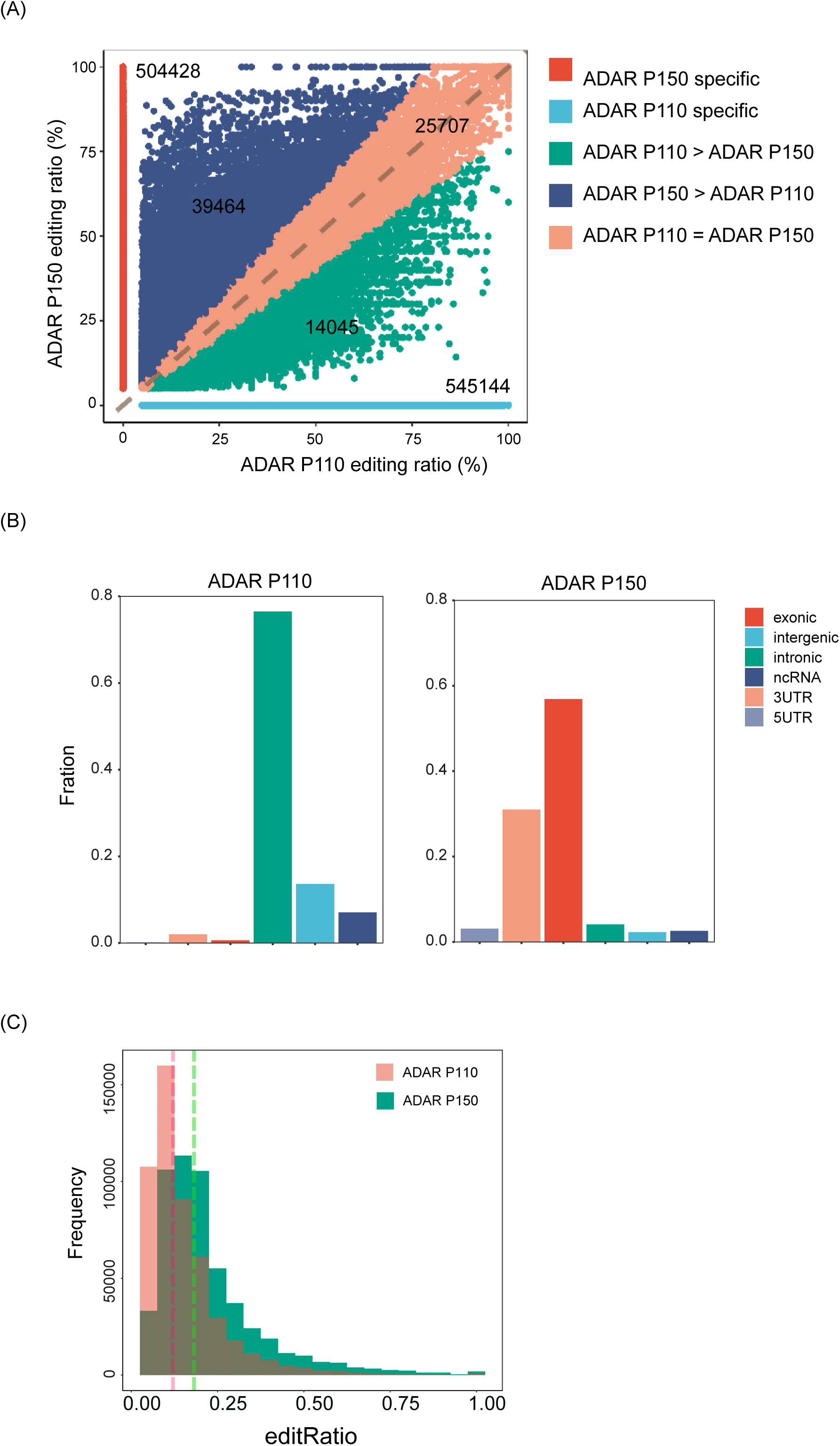
Z-α domain protein (ADAR1 P150) preferentially localizes to RNA exonic and 3’UTR regions. (A) A scatter plot comparing the distribution of A-to-I editing sites in cells overexpressing ADAR1 P110 versus ADAR1 P150. This visualization highlights the distinct editing site preferences between the two isoforms. (B) Detailed analysis of the localization of A-to-I editing sites. In cells overexpressing ADAR1 P110, editing sites are primarily found in intronic regions (depicted in green). In contrast, cells overexpressing ADAR1 P150 show a notable enrichment of A-to-I editing sites in exonic (depicted in red) and 3’UTR (depicted in pink) regions, demonstrating the unique targeting of Z-α Domain Protein (ADAR1 P150) to these specific transcriptomic areas. (C) Histogram displaying the frequency distribution of A-to-I editing efficiency in cells overexpressing either ADAR1 P110 or ADAR1 P150. This analysis reveals differences in the median editing efficiency between the two groups, underscoring the distinct editing preferences of RNA substates associated with each ADAR1 isoform.

